# Multi-omics integration identifies a reproducible inflammatory host-response axis in pediatric sepsis

**DOI:** 10.64898/2026.07.06.736269

**Authors:** Mariam Ait Oumelloul, Ali Saadat, Vito Zanotelli, Barry Ryan, Dylan Lawless, Andrea Agostini, Simon Tang, Daphné Chopard, Ivana Nemes-Bokun, Victoria J. Wright, Jethro Herberg, Christa E. van der Gaast-de Jongh, Marien de Jonge, Vanessa Sancho-Shimizu, Mike Levin, Philipp K.A. Agyeman, Christoph Berger, Nicola Zamboni, Sandra Goetze, Cédric Howald, Katrin Männik, Rebeca Mozun, Luregn J. Schlapbach, D. Sean Froese, Jacques Fellay, Swiss Pediatric Sepsis Study, EUCLIDS Consortium, SwissPedHealth Consortium

**Author notes:** These authors jointly supervised this work (Luregn J. Schlapbach, D. Sean Froese, Jacques Fellay).

## Abstract

Sepsis is a major cause of morbidity and mortality in children, yet biological heterogeneity in host responses has limited progress toward targeted therapies and patient stratification. Multiomics integration can combine complementary molecular layers to identify coordinated disease programs not captured by individual assays. Here, we integrated genomic, bulk transcriptomic, proteomic and metabolomic data from blood samples of 22 children with culture-confirmed bacterial sepsis enrolled in the Swiss Pediatric Sepsis Study. Multi-Omics Factor Analysis identified a dominant host-response axis reflecting systemic inflammation. This axis was driven primarily by transcriptomic variation and supported by coordinated proteomic and metabolomic signals, including circulating inflammatory mediators and altered amino-acid metabolism. It was associated with C-reactive protein and a severity score proxy. Projection into an independent pediatric sepsis cohort (n = 22) reproduced the inflammatory and severity-related interpretation of this axis. Single-omic projections showed that the integrated signal could be approximated from individual layers, particularly transcriptomics. In three external pediatric whole-blood transcriptomic datasets, the RNA-derived projection separated septic shock from healthy controls and increased across clinical inflammatory syndromes. These findings define a reproducible inflammatory host-response axis in pediatric sepsis and support multi-omics-guided selection of molecular readouts suitable for clinical translation.

## 1 INTRODUCTION

Sepsis, defined as a dysregulated host response to infection leading to life-threatening organ dysfunction, affects approximately 50 million people globally each year, half of whom are neonates and children under 19 years^1^. This disproportionate pediatric burden contributes substantially to global child mortality, morbidity, and developmental consequences^2^. Despite decades of research, therapeutic advances have been limited^3,4^, in part because sepsis represents a biologically heterogeneous syndrome rather than a single disease entity. This heterogeneity complicates diagnosis, prognosis, and the development of targeted interventions^4^.

Systems biology approaches have begun to uncover distinct molecular patterns underlying host responses to infection. Transcriptomic studies, in particular, have identified gene-expression signatures that stratify patients by immune state and clinical outcome. For instance, one framework delineated two core transcriptional states, resistance and systemic inflammation, whose relative balance reflects disease severity, with sepsis characterized by a diminished resistance-to-inflammation ratio^5^.

Similarly, the Sepsis Response Signature (SRS) framework defined transcriptional states associated with immune suppression and differential disease severity, classifying patients into discrete immune states (SRS1, SRS2, and SRS3) based on 7- or 19-gene signatures^6,7^. Individuals with an SRS1 profile exhibit marked immune suppression, greater organ dysfunction, and higher mortality than those with an SRS2 profile. These transcriptional states have also been linked to cellular immune phenotypes, including immature neutrophil expansion and altered adaptive immune responses^8^. Building on this classification, a continuous severity score termed SRSq was later introduced to model disease as a spectrum rather than as discrete states. Notably, SRSq was further validated in pediatric bacterial and viral sepsis, where it captured an acute illness signal and discriminated between degrees of disease severity^7^.

However, transcriptomic signals alone do not fully capture the biology of sepsis, partly because RNA abundance is shaped by post-transcriptional regulation and does not directly reflect circulating proteins, metabolic states, or downstream effector processes. Consistent with this, proteomic studies of sepsis have revealed complementary patterns of immune activation and metabolic disruption that only partially overlap with transcriptional signatures^9^. Integrating multiple molecular layers may therefore enable the identification of molecular signatures analogous to SRS while extending beyond transcriptome-centred approaches by capturing complementary biological information and, potentially, more accessible surrogate biomarkers.

In the present study, we performed integrated genomic, bulk transcriptomic, proteomic, and metabolomic profiling of blood samples from children with culture-confirmed bacterial sepsis (Figure 1). Using Multi-Omics Factor Analysis (MOFA)^10^, we identified latent molecular factors capturing shared variation across omics layers and evaluated their biological and clinical relevance. By projecting these factors into independent cohorts and external transcriptomic datasets, we assessed the robustness and generalizability of the identified host-response programs.

**Figure 1:**
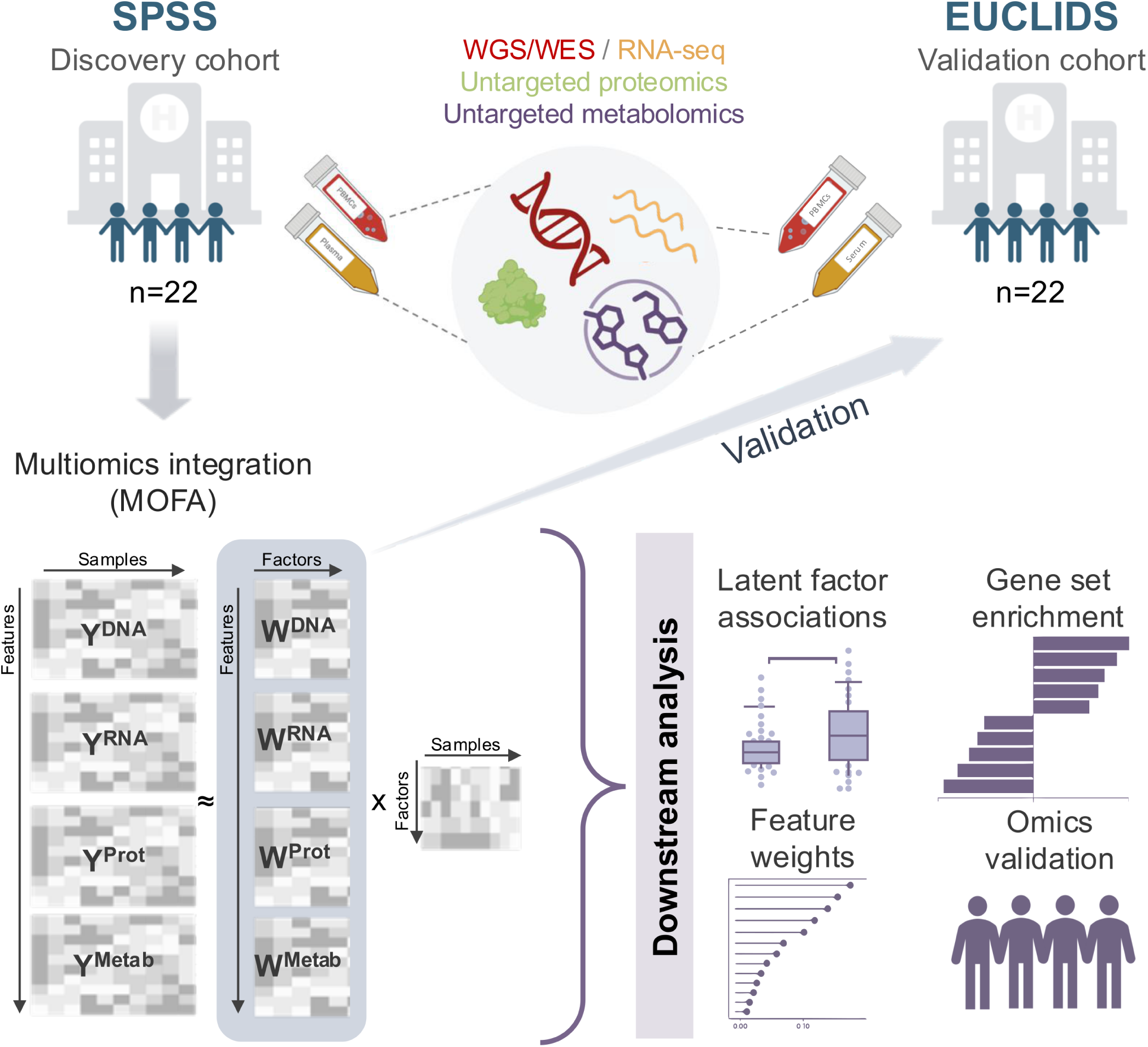
Study design and multi-omics integration workflow. Two independent cohorts were analysed: SPSS (discovery cohort; n = 22) and EUCLIDS (validation cohort; n = 22). For each cohort, blood samples were profiled using whole-genome/whole-exome sequencing (WGS/WES), RNA sequencing (RNA-seq), untargeted proteomics and untargeted metabolomics. Multi-omics layers from the discovery cohort were integrated using Multi-Omics Factor Analysis (MOFA) to infer latent factors capturing shared and modality-specific sources of variation across datasets. Factors were carried forward for downstream analyses, including association of latent factors with clinical or biological variables, identification of key contributing features via factor weights, gene set enrichment of factor-associated gene programs, and orthogonal validation of omics signals. Findings were evaluated in the independent EUCLIDS cohort to assess robustness and reproducibility of multi-omics-derived factors and associated signatures. SPSS, Swiss Pediatric Sepsis Study; EUCLIDS, European Union Childhood Life-threatening Infectious Disease Study.

## 2 RESULTS

### 2.1 Comprehensive multi-omics profiling of pediatric sepsis

We performed multi-omics profiling in pediatric bacterial sepsis patients from the SPSS cohort (n = 22) and validated the key biological signatures in an independent sub-cohort from EUCLIDS (n = 22) (Figure 1).

The SPSS cohort included a higher proportion of males than EUCLIDS (68% vs 50%), whereas EUCLIDS participants were younger on average (median age 485.5 days vs 2,589.5 days in SPSS). All EUCLIDS patients were admitted to the pediatric intensive care unit (PICU), while intensive care was required for only a minority of SPSS cases (18%). Only a few deaths were reported in this study, occurring exclusively among patients in the EUCLIDS cohort (n = 4). Mortality was therefore not analyzed as an outcome variable, recognizing that the number of fatal cases was too small to allow meaningful statistical inference (Table 1).

**Table 1:**
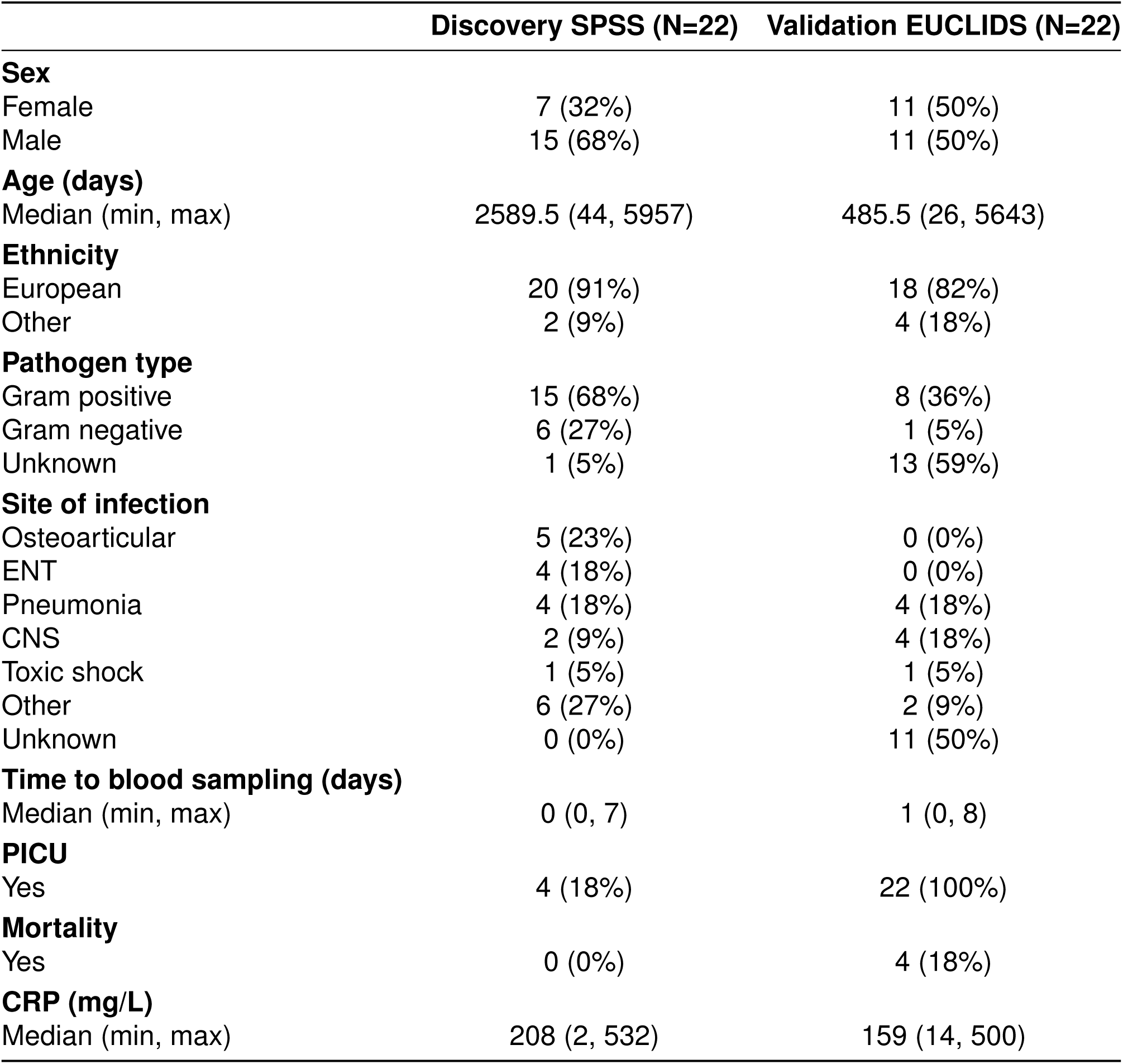
Comparison of Discovery (SPSS) and Validation (EUCLIDS) cohorts.

Multi-omics data were generated across four molecular layers for both cohorts: genomics (WGS/W bulk transcriptomics from whole blood, untargeted proteomics and metabolomics (Supplementary Figure S2). Proteomic and metabolomic profiling were performed on serum samples in SPSS and on plasma samples in EUCLIDS.

### 2.2 Systematic feature selection defines robust multi-omic latent axes

To identify shared molecular programs underlying pediatric sepsis across multiple data modalities, we applied MOFA framework^10^ to the discovery SPSS cohort.

We performed a grid search to optimize the number of input features for DNA, RNA, and RNA splicing by evaluating different values of N corresponding to the most variable features in each layer. This strategy aimed to maximize the variance explained by the model and capture the most informative representation of biological signal. The variance explained was consistently dominated by the transcriptomic layer, and increasing the number of RNA features beyond 1,500 yielded minimal gains in total variance while disproportionately reducing the contributions of proteomics and metabolomics (Supplementary Figure S3A). In contrast, DNA contributed minimally to overall variance in this model, consistent with sepsis being largely environment and response-driven rather than genetically determined. However, excluding the DNA view (DNA = 0) did not enhance model performance, confirming that while DNA accounted for a small fraction of variance, retaining it contributed to a more comprehensive multi-omic representation (Supplementary Figure S3B). Based on the grid search results, we selected 1,500 RNA, 500 RNA splicing, and 500 DNA features as the optimal configuration for downstream modeling (highlighted with “X” in Supplementary Figure S3B). For proteomics and metabolomics, we retained all features detected by the respective technologies, resulting in 637 proteomic and 1,943 metabolomic features.

After fixing the number of features, we assessed latent factors stability. Across 25 independent MOFA runs initialized with different random seeds, the inferred latent structure was highly consistent, with strong pairwise correlations between matched components across runs, indicating robust and reproducible model inference (Supplementary Figure S4).

### 2.3 Variance explained across omics layers

After feature selection, we trained the MOFA model using the 22 pediatric sepsis samples from the SPSS cohort (Figure 2A). The inferred latent factors captured both shared sources of variation across data types and modality-specific patterns. Overall, the variance explained was largely driven by RNA (71.71%), indicating that gene expression accounts for most of the variability across samples captured by the model. Proteomics and metabolomics also made meaningful contributions across multiple factors, with total variance explained of 51.00% and 33.70%, respectively (Figure 2B).

**Figure 2:**
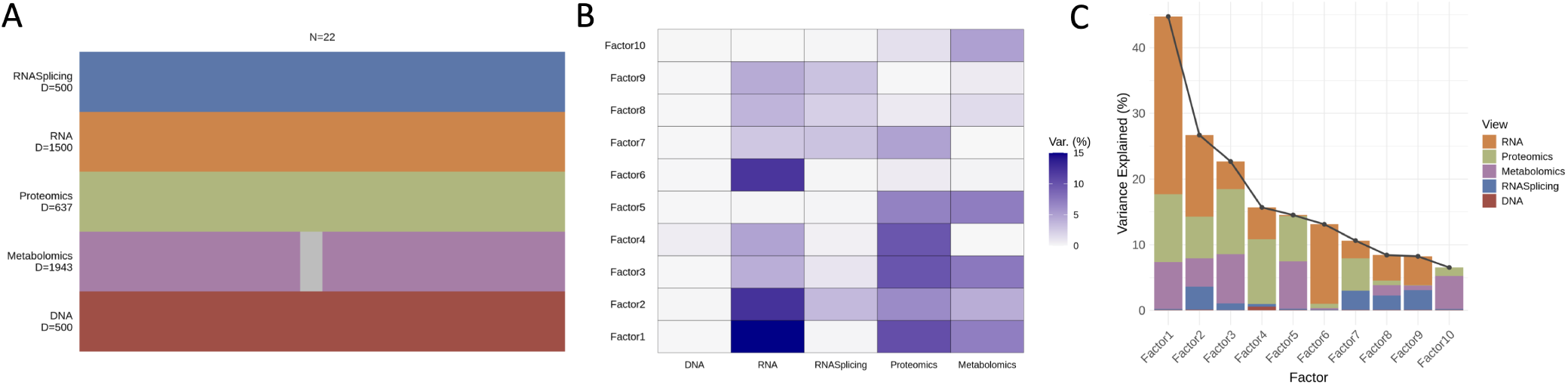
Multi-omics integration with MOFA reveals shared axes of biological variation. (A) Overview of data modalities included in the MOFA model. Data modalities are shown in different rows (D = number of features) and samples (N) in columns, with missing samples shown using grey bars. (B) Fraction of variance explained by each omic layer across the ten inferred latent factors. (C) Total variance explained by each factor per modality.

Cumulative variance plots indicated that the first three factors captured most of the signal across omics layers (Figure 2C). Factor 1, which represents the dominant axis of variation, was primarily driven by transcriptomics (27.03% variance explained), with additional contributions from proteomics (10.34%) and metabolomics (7.15%). Factor 2 reflected a secondary orthogonal axis, characterized by coordinated molecular changes across transcriptomics (12.39%), proteomics (6.37%), metabolomics (4.29%), and RNA splicing (3.53%).

### 2.4 Statistical associations between latent factors and demographic and clinical variables

We next assessed associations between latent factors and available demographic and clinical variables (Figure 3A). Factor 1 values were positively and significantly correlated with C-reactive protein (CRP), the most commonly used clinical marker of inflammation, measured by routine blood testing (Spearman *ρ*= 0.51, p=1.5e-3) (Figure 3B) and with proteomics-derived CRP (Spearman *ρ*=0.62, P=2e-3) (Figure 3C), indicating concordance between clinical and molecular readouts of systemic inflammation. Factor 1 also showed a strong positive association with SRSq, a quantitative score reflective of degree of immune dysfunction, (Spearman *ρ*=0.93, P=4.10e-6), further linking factor 1 to clinical severity. In line with this, the highest factor 1 score was observed in a patient diagnosed with septic shock (patient S4, Figure 3 A-C). Factor 2 was associated with patient age, with the lowest scores in children *<* 4 years and progressively higher values in older patients, consistent with an age-dependent latent program. (Spearman *ρ*=0.57, P=5.6e-3) (Figure 3 D). This pattern suggests developmental regulation of the underlying molecular processes, distinct from the inflammation–severity axis captured by factor 1.

**Figure 3:**
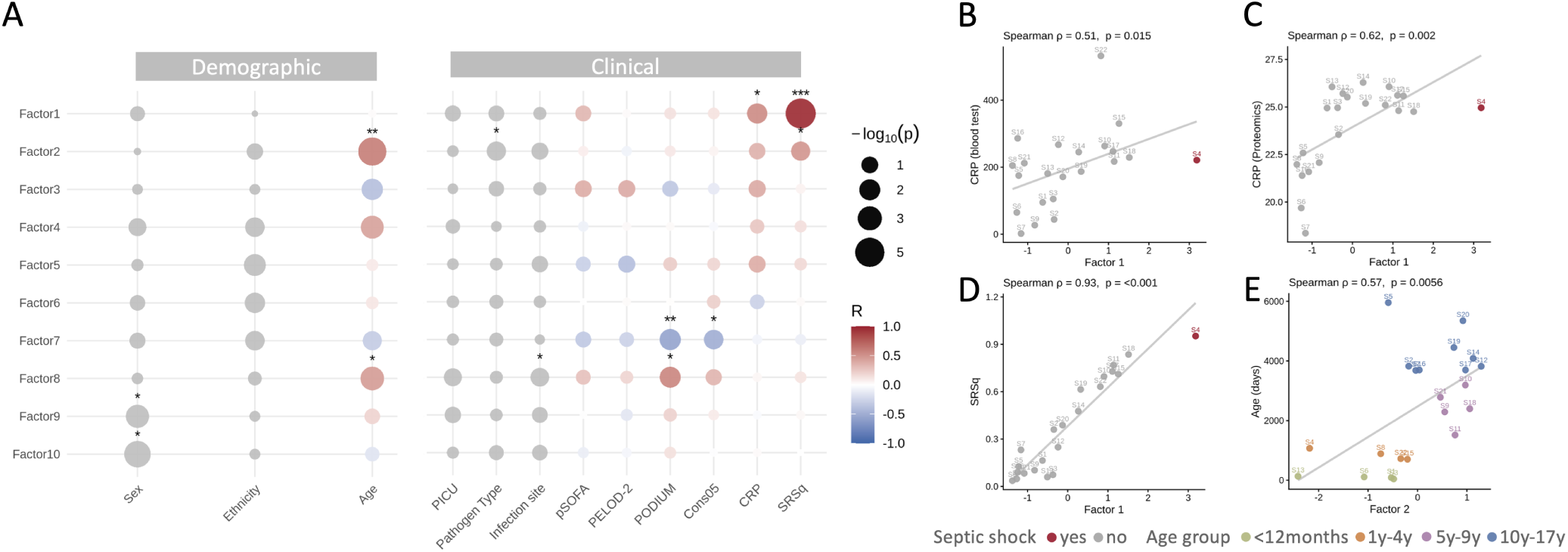
Clinical and demographic correlates of MOFA latent factors in pediatric sepsis. (A) Bubble plot summarizing associations between each MOFA factor and demographic or clinical variables. Circle color indicates the direction and magnitude of the Spearman correlation coefficient for continuous variables (red: positive, blue: negative), and circle size reflects statistical significance (log_10_(P) with stars indicating significance levels (∗, ∗∗, ∗ ∗ ∗ denote P *<* 0.05, 0.01, and 0.001, respectively). (B-D) Relationships between Factor 1 scores and CRP measured by clinical blood testing (B), CRP quantified in the proteomic layer (C), and severity score (SRSq) (D). Septic shock status is indicated (red, yes; grey, no). (E) Relationship between Factor 2 scores and age (days). Points are colored by age group (*<*12 months, 1–4 years, 5–9 years, 10–17 years). PICU, pediatric intensive care unit; pSOFA, pediatric Sequential Organ Failure Assessment; PELOD-2, pediatric Logistic Organ Dysfunction-2; PODIUM, pediatric Organ Dysfunction Information Update Mandate; Cons05, 2005 consensus score; CRP, C-reactive protein; SRSq, Quantitative sepsis response signature

Factors 7 and 8 showed moderate correlations with the Pediatric Organ Dysfunction Information Update Mandate (PODIUM) score (Spearman *ρ*=-0.54 P=8.87e-3 and Spearman *ρ*=0.54 P=1.01e-2, respectively) (Supplementary Figure S5). No significant associations were observed with other clinical severity scores, including pSOFA, or with PICU admission status.

### 2.5 Inflammatory axis dominates cross-omics molecular variation in pediatric sepsis

Among the latent factors identified by MOFA, factor 1 captured the dominant axis of biological variation and reflected a coherent multi-omic program of systemic inflammation. Factor 1 was primarily driven by transcriptomics, with top positive loadings including neutrophil-associated genes (e.g. *CD177*, *MMP8*, *CHIT1*) and enrichment for neutrophil degranulation and innate immune pathways (Figure 4 A-B). Concordantly, correlation of CIBERSORT-derived immune-cell estimates with MOFA scores linked factor 1 to increased relative neutrophil abundance (Spearman *ρ* = 0.79, P = 1.60e-5) and reduced adaptive immune-cell proportions, particularly CD8^+^ T cells (Spearman *ρ* = −0.90, P = 1.03e-8) (Supplementary Figure S6). Proteomic loadings provided complementary information, with acute-phase mediators positively weighted (e.g. *PTX3*, *SAA1*/*SAA2*) (Figure 4C). Enrichment analyses implicated stress-response and apoptosis-related processes (Figure 4D). Metabolomic negative loadings mapped to chemical formulas consistent with amino acids (including proline, tyrosine, tryptophan and citrulline), suggesting amino-acid depletion during inflammatory stress (Figure 4E). Unsupervised clustering of top features across layers stratified samples by inflammatory status and recapitulated gradients in blood CRP and SRSq, supporting factor 1 as a multi-omic correlate of inflammatory burden and severity.

**Figure 4:**
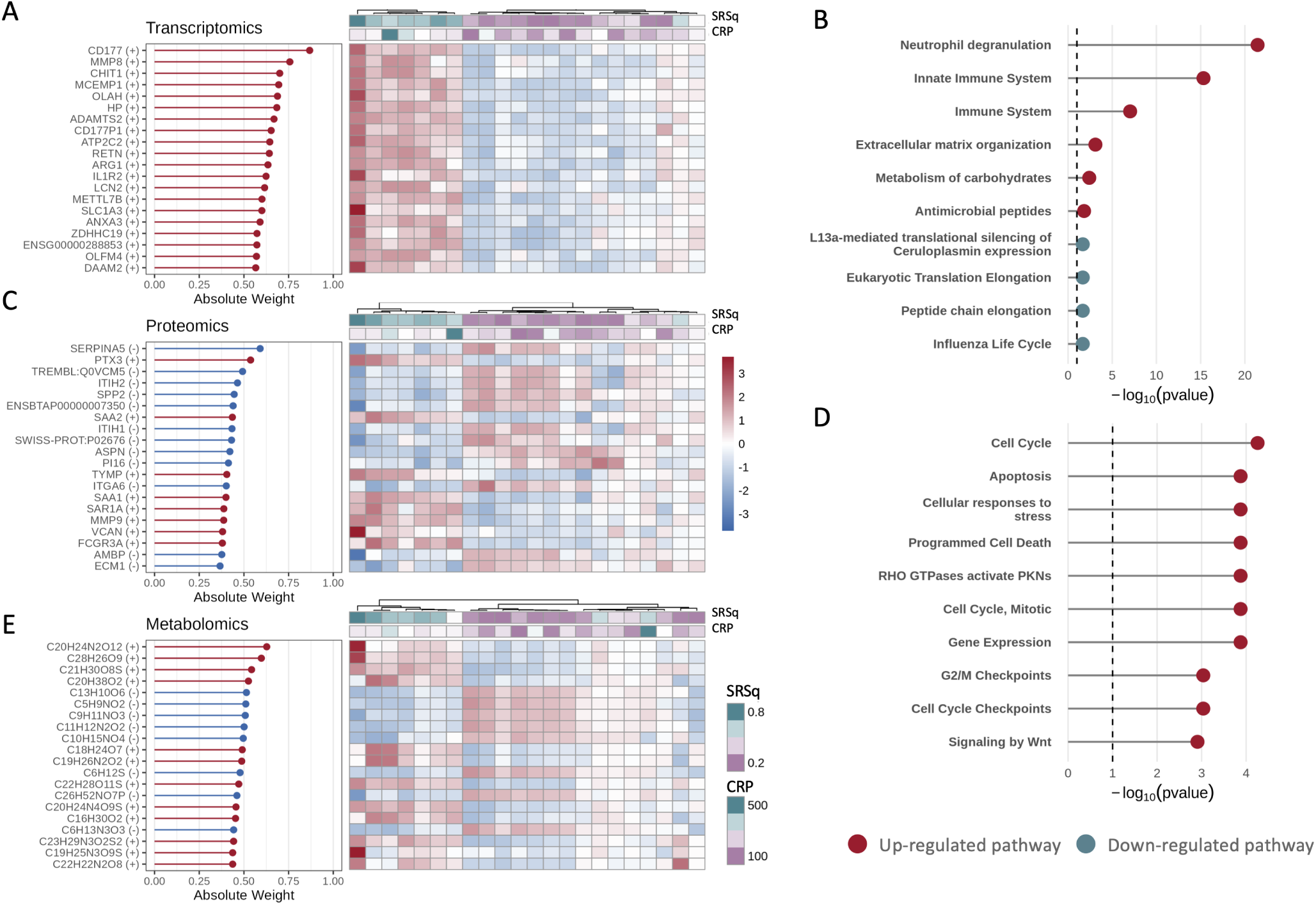
Factor 1 defines a coordinated inflammatory multi-omic axis. (A) Top 20 features with the highest absolute weights contributing to the first latent factor across transcriptomic dataset. Their normalized expression are displayed as heat maps (right) where samples are ordered by unsupervised hierarchical clustering of columns using the Ward.D2 method, and annotation bars denote SRSq and blood CRP levels. (B) Gene set enrichment analysis (GSEA) of pathways associated with transcriptomics latent factor weights. (C) Features with the highest absolute weights contributing to the first latent factor across proteomics dataset. (D) Gene set enrichment analysis (GSEA) of pathways associated with proteomics latent factor weights. (E) Features with the highest absolute weights contributing to the first latent factor across metabolomics dataset. The top 20 features per layer are shown, ranked by absolute weight (left), and their normalized expression or abundance profiles are displayed as heat maps (right). Samples are ordered by unsupervised hierarchical clustering using the Ward.D2 method, and annotation bars denote SRSq and CRP levels.

### 2.6 Multi-omic validation of factor 1 in the independent EUCLIDS cohort

To assess the robustness and generalizability of factor 1, we projected the MOFA-derived weights into an independent pediatric sepsis cohort from EUCLIDS. In this cohort, proteomic and metabolo data were generated from plasma, in contrast to the serum-based assays in the discovery (SPSS) cohort. Despite these matrix differences, factor 1 preserved its inflammatory interpretability and showed a moderate correlation with clinically measured CRP (Spearman *ρ* = 0.49, P = 5.4e-2), proteomics-derived CRP abundance (Spearman *ρ*=0.53, P=1.7e-2) and a strong correlation with the SRSq score (Spearman *ρ* = 0.89, P = 2.6e-5) (Figure 5 A-C). Consistent with observations in the discovery cohort, CIBERSORT-based immune-cell deconvolution linked higher factor 1 scores to increased relative neutrophil abundance (Spearman *ρ* = 0.66, *P* = 1.29×10*^−^*^2^) and reduced CD8^+^ T cells (Spearman *ρ* = −0.59, *P* = 3.03×10*^−^*^2^) (Supplementary Figure S7).

**Figure 5:**
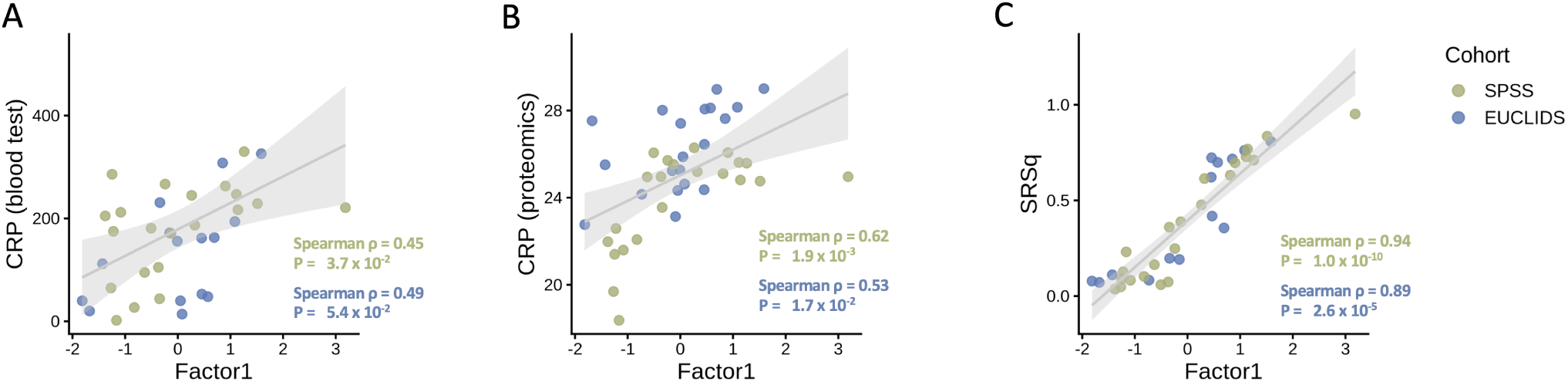
The inflammatory factor is reproduced in an independent pediatric sepsis cohort. Correlation between factor 1 and clinical or molecular severity measures across the EUCLIDS (blue) and SPSS (green) cohorts. Each point represents an individual patient, and dashed lines denote linear regression fits for each cohort. The Spearman correlation coefficient (R) and corresponding P value are shown per cohort. (A) C-reactive protein (CRP) blood levels. (B) CRP intensity from proteomics data. (C) Sepsis Response Signature (SRSq) score.

### 2.7 Single-omic projections recover the clinical inflammatory axis captured by factor 1

Because factor 1 captured a clinically relevant axis of inflammation and severity, defined by its association with CRP and SRSq, we next asked whether this multi-omic signal could be approximated using individual omic layers alone. This analysis was motivated by the potential translational value of identifying single-modality proxies in settings where full multi-omics profiling is not feasible. In the discovery SPSS cohort, projections based on RNA, proteomics and metabolomics each retained positive associations with CRP (RNA: Spearman *ρ* = 0.53, *P* = 0.011; proteomics: *ρ* = 0.57, *P* = 5.6 × 10*^−^*^3^; metabolomics: *ρ* = 0.56, *P* = 8.1 × 10*^−^*^3^; Figure 6A-C). Associations with CRP were weaker in the independent EUCLIDS cohort (Spearman RNA: *ρ* = 0.49, *P* = 0.086; proteomics: *ρ* = 0.40, *P* = 0.121; metabolomics: *ρ* = 0.42, *P* = 0.139; Figure 6A-C), potentially reflecting differences in sample matrix, as proteomic and metabolomic profiling was performed in serum in SPSS and plasma in EUCLIDS. Consistent with the clinical relevance of factor 1, RNA-, proteomics- and metabolomics-based projections were also positively associated with SRSq in SPSS (RNA: *ρ* = 0.91, *P <* 10*^−^*^4^; proteomics: *ρ* = 0.75, *P <* 10*^−^*^4^; metabolomics: *ρ* = 0.85, *P <* 10*^−^*^4^; Figure 6 D-F). In EUCLIDS, RNA and metabolomics retained strong associations with SRSq (RNA: Spearman *ρ* = 0.93, *P <* 10*^−^*^4^; metabolomics: *ρ* = 0.80, *P* = 0.0019), whereas the proteomics-based projection showed a more modest trend (proteomics: *ρ* = 0.42, *P* = 0.13; Figure 6 D-F). Together, these results indicate that the inflammatory severity signal captured by factor 1 can be approximated from individual omic layers, with RNA and metabolomics providing the most consistent single-layer proxies of the integrated multi-omic axis in both datasets.

**Figure 6:**
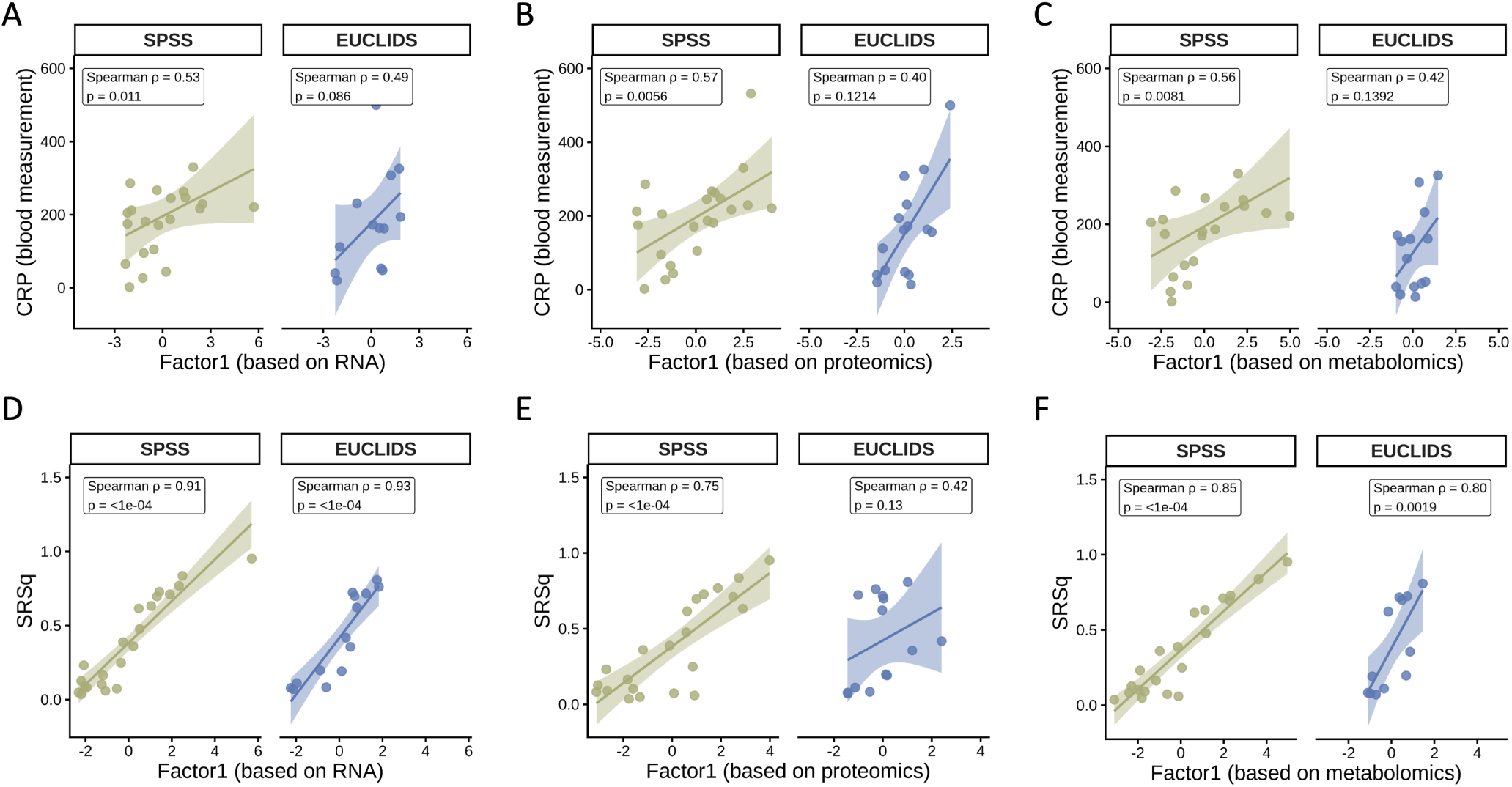
Single-omic projections recover the inflammatory severity axis captured by factor 1. Associations between factor 1 projections derived from individual omic layers and clinical measures of inflammation and severity in the SPSS discovery cohort and the independent EUCLIDS cohort. (A-C) Associations between CRP measured in blood and factor 1 projections based on (A) RNA, (B) proteomics and (C) metabolomics. (D-F) Associations between SRSq and factor 1 projections based on (D) RNA, (E) proteomics and (F) metabolomics. Points represent individual participants. Lines show fitted linear trends with shaded 95% confidence intervals. Spearman correlation coefficients and two-sided P values are shown in each panel.

### 2.8 Transcriptomic projections validate factor 1 as a severity-associated inflammatory axis

Having established that factor 1 can be approximated from individual omic layers, with RNA providing one of the most consistent single-modality proxies, we next asked whether its transcriptomic component was reproducible across larger independent pediatric sepsis cohorts. We therefore projected the factor 1 gene-weight vector onto three publicly available transcriptomic datasets (GSE26440, GSE26378 and GSE13904), each including healthy controls. Factor 1 was consistently positively correlated with SRSq across all three cohorts (Spearman *ρ*= 0.86, P = 1.06e-39 for GSE26440; Spearman *ρ*= 0.86, P = 7.44e-32 for GSE26378; and Spearman *ρ*= 0.84, P= 1.52e-37 for GSE13904 (Supplementary Figure S8). Across all three cohorts, Factor 1 distinguished septic shock patients from healthy controls (Wilcoxon test, P *<* 0.001), with controls exhibiting consistently negative scores (Figure 7 A-B). In GSE13904, Factor 1 values increased progressively across the ordered clinical spectrum from controls to SIRS, sepsis and septic shock, with median scores highest in septic shock, followed by sepsis and SIRS. Consistent with this pattern, cumulative link ordinal regression showed that Factor 1 was significantly associated with increasing disease severity across the clinical spectrum (all P *<* 0.001; Figure 7 C), suggesting that this axis may reflect a continuum of inflammatory activation rather than sepsis-specific transcriptional programs. Together, these cross-cohort validations demonstrate that Factor 1, the principal source of heterogeneity within the cohort, constitutes a robust and biologically reproducible axis of inflammation that scales with disease severity, and is consistently observed across independent datasets.

**Figure 7:**
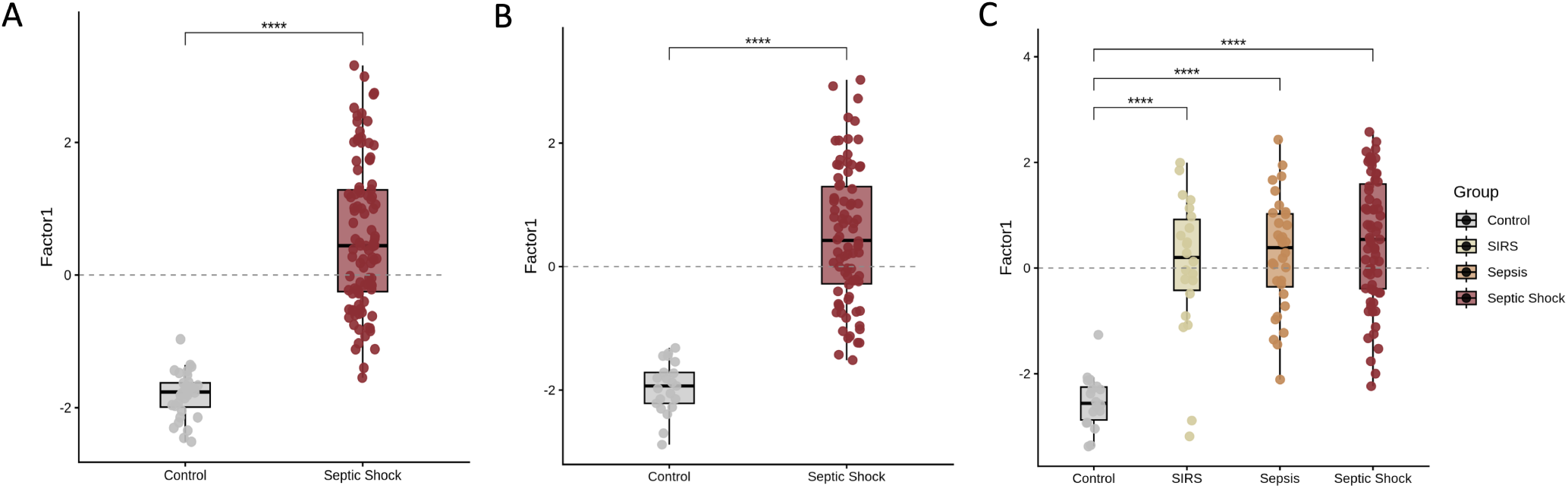
Validation of factor 1 across multiple independent transcriptomic datasets. Each boxplot compares factor1 scores between healthy controls and sepsis-related clinical groups; center lines indicate medians, boxes represent interquartile ranges, whiskers indicate 1.5× the interquartile range, and points denote individual samples. Statistical significance was assessed using Wilcoxon rank-sum tests (**** P *<* 0.0001). (A) GSE26378 pediatric cohort . (B) GSE26440 pediatric cohort. (C) GSE13904 pediatric cohort showing control, SIRS, sepsis, and septic shock groups.

## 3 DISCUSSION

Our results reveal a reproducible inflammatory axis that integrates signals across molecular layers and associates with disease severity in pediatric sepsis. Given the marked heterogeneity of host responses in sepsis, this axis provides a framework for capturing clinically relevant variation across patients beyond transcriptomic signatures alone. Although driven primarily by transcriptomic variation, it was reinforced by coordinated proteomic and metabolomic changes, linking cellular immune activation to systemic inflammatory and metabolic responses. This cross-omic structure highlights complementary biological pathways that could support more refined patient stratification and enable the development of targeted, pragmatic biomarker panels. Importantly, the inflammatory program was reproducible across independent pediatric cohorts and across molecular platforms, demonstrating that it represents a robust feature of the host response to severe infection rather than a cohort-specific signal.

The first latent factor identified in our analysis (factor 1) correlated with blood CRP levels and with the SRSq score, a measure of sepsis severity^7^. Although SRSq is constructed from a 19-gene transcriptomic signature, the features loading onto factor 1 showed only limited direct overlap with these genes (N=2, *TDRD9* and *ZAP70*).

Mechanistically, factor 1 is dominated by a neutrophil-driven innate immune transcriptomic program, consistent with a hyperinflammatory host-response state characterized by enhanced degranulation and broad pro-inflammatory activation. Dysregulated neutrophil activation is a recognised driver of tissue injury and adverse outcomes in sepsis, linking this transcriptional pattern to clinically meaningful severity^11^. The strong contribution of *CD177*, a neutrophil activation marker, is in line with prior reports describing high *CD177* as a hallmark of severe disease^12^. *MMP8*, an endopeptidase predominantly produced by neutrophils, supports neutrophil infiltration and effector function^13^, and together with *CHIT1*^14^ points to a neutrophil-associated inflammatory axis. Notably, the presence of *IL1R2* and *ARG1* among the top-loading features further suggests a shift toward immature neutrophil programs, as these markers are consistently upregulated in immature neutrophil populations in single-cell studies of sepsis and have been linked to emergency granulopoiesis and immunosuppressive neutrophil states^8^.

Consistent with this neutrophil-associated loading structure, CIBERSORT-based deconvolution linked higher factor 1 scores to increased neutrophil proportions, reinforcing its interpretation as an innate inflammatory axis. The same analysis also identified a marked negative association with CD8^+^ T-cell abundance, suggesting that this neutrophil-dominated programme may be coupled to depletion of adaptive immune compartments. This pattern is concordant with single-cell studies of sepsis showing expansion of immature and emergency-granulopoiesis neutrophil states alongside lymphocyte depletion, suggesting that factor 1 captures a bulk multi-omic representation of immune-cell remodelling during infection^8,15^.

The proteomic signature reinforces this same programme through increased abundance of myeloid and neutrophil-associated proteins, including *MMP9* (granule-derived protease) and *FCGR3A*, consistent with downstream effector release. At the same time, proteomics provides information that is less accessible from transcriptomics alone by directly capturing the circulating inflammatory milieu: elevated acute-phase and soluble innate mediators (*SAA1/SAA2*, *PTX3*) reflect systemic inflammation, whereas reduced serine protease inhibitors (*SERPINA5* and *ITIH1/ITIH2*) are compatible with down-regulation of protective protease and coagulation regulators during severe disease. Metabolomics further complements these layers by associating factor 1 with broad amino-acid depletion (e.g., citrulline, tryptophan, tyrosine and proline), consistent with infection-driven reprogramming of host amino-acid metabolism. Overall, this integrated multi-omic view illustrates how transcriptomics captures the cellular activation program, while proteomics and metabolomics read out downstream effector release and organism-wide host-response changes in the circulation.

Although the axis was identified using multi-omic integration, it could be recovered using single-omic projections: factor 1 derived from transcriptomics-, proteomics-, or metabolomics-only data retained consistent associations with CRP and SRSq in both discovery and validation cohorts. This supports a practical discovery-to-deployment workflow in which comprehensive multi-omic profiling is used upfront to define latent axes that integrate signals across molecular layers and provide a mechanistic interpretation, after which a single, clinically tractable assay can be selected based on how faithfully each layer reconstructs the axis and how readily it can be measured in routine practice.

Given that factor 1 is largely driven by transcriptomic features, and that a transcriptomics-only projection retained its associations with inflammation and clinical severity, we next evaluated its behavior in larger external pediatric datasets that included controls. Factor 1 scores were substantially lower in controls, yielding clear separation from septic shock cases. Factor 1 also separated SIRS from controls and arranged SIRS, sepsis and septic shock along a graded trajectory, suggesting an inflammatory continuum rather than discrete categories. By contrast, Factor 1 did not consistently discriminate survivors from non-survivors (Supplementary Figure S9), indicating that this hyperinflammation axis captures severity but is not sufficient to explain prognosis, and motivating the contribution of additional latent factors.

Factor 2 showed a consistent positive association with age at blood collection in both cohorts (SPSS: R = 0.57, P = 6.5e-3; EUCLIDS: R = 0.65, P = 1.9e-3; Supplementary Figure S10), identifying a reproducible age-related molecular axis. As MOFA latent factors capture orthogonal axes of variation, the separation between factor 1 and factor 2 suggests that age-related molecular variation represents an additional dimension of host heterogeneity beyond the hyperinflammatory severity programme captured by factor 1. Across molecular layers, the features contributing to factor 2 converge on extracellular matrix organisation (*TGFBI*, *HTRA1*, *COL12A1*), complement activity (*C9*, *CFHR4*) and coagulation pathways (*F9*), supported by enrichment of matri-some and complement/coagulation processes (Supplementary Figure S11). Similar pathways have consistently been identified as major determinants of age-related variation in the circulating proteome, particularly during childhood^16,17^, supporting the interpretation that factor 2 captures developmental and homeostatic biology distinct from the acute inflammatory programme represented by factor 1.

This study has several limitations. First, the sample size was modest and comprehensive pediatric sepsis multi-omics datasets remain scarce, which constrained statistical power and limited opportunities for external validation in larger cohorts beyond transcriptomics. Second, clinical heterogeneity across cohorts complicated direct clinical anchoring: pediatric sepsis severity is captured by multiple scoring frameworks that are not fully harmonized across studies, and mortality events were too few to robustly model hard outcomes, restricting inference about prognostic performance beyond severity-associated correlates. Finally, while we integrated genomics and splicing signals, alternative integration strategies may better capture their distinct biology. In addition, splicing-based signals were less intuitive to interpret than expression changes.

Future work should prioritize projection of the learned factor weights into larger, independent pediatric multi-omics cohorts as such resources emerge, enabling more precise estimation of factor stability, subgroup structure and cross-platform transferability. To facilitate this, we share the learned factor weights with the community for independent replication and benchmarking (Supplementary Table ST1-ST3). In parallel, linking factor scores to richer and more standardised clinical metadata, including longitudinal trajectories, organ dysfunction phenotypes, treatment exposures and outcomes, will be essential to distinguish correlates of acute severity from predictors of prognosis. While inflammatory severity appears well captured by transcriptomic-dominant programs, it remains unresolved whether integrated multi-omic factors improve prediction of mortality or other hard endpoints, or instead primarily refine mechanistic stratification that can guide host-directed interventions. Addressing these questions will require larger studies with standardised sampling windows, harmonised clinical scoring and sufficient outcome events.

## 4 METHODS

### 4.1 Study Cohort

This study was conducted as part of the Lighthouse project of the SwissPedHealth national data stream initiative, a nationwide effort to make routine clinical data from Swiss pediatric hospitals standardized, interoperable, and usable for research^18^. Children aged ≤ 17 years with blood culture–confirmed bacterial sepsis were eligible for inclusion. Patients were retrospectively selected from two independent cohorts, used for discovery and validation, respectively: the Swiss Pediatric Sepsis Study (SPSS)^19^ and the European Union Childhood Life-threatening Infectious Disease Study (EUCLIDS)^20^. The SPSS is a national, multicenter, observational cohort study investigating blood culture–proven bacterial sepsis in children across Switzerland. EUCLIDS is a multicenter cohort study enrolling children presenting to hospitals with life-threatening bacterial infections. Comprehensive clinical and laboratory data were collected and harmonized across both cohorts. For each participant, a frozen aliquot of biological material was used for whole-genome sequencing (WGS), RNA sequencing (RNA-seq), and data-independent acquisition mass spectrometry (DIA–MS). Where available, frozen aliquots of biological material were used for whole-genome sequencing (WGS), RNA sequencing (RNA-seq), protein and metabolite profiling. For some participants, whole-exome sequencing (WES) data generated previously were also included^21^.

The SPSS study was approved by the Ethics Committee of the Canton of Bern (approval number KEK-Nr. 029/2011). The reuse of the samples from SPSS for the present analysis was approved by the Ethics Committee of the Canton of Zurich using an exemption according to Art. 34 HRA (BASEC 2022-00351). The EUCLIDS study protocol was approved by the United Kingdom Research Ethics Committee (approval number 11/LO/1982). Only samples for which appropriate ethical approval and informed consent were obtained were included in this analysis.

### 4.2 Multi-omics measurements

#### 4.2.1 Genomics

WGS was performed for 24 individuals, including 7 from the SPSS cohort and 17 from the EUCLIDS cohort. For an additional 15 SPSS samples, previously generated whole-exome sequencing (WES) data were available, and the corresponding raw FASTQ files were incorporated into the analysis^21^. For WGS, genomic DNA was isolated from whole blood. Sequencing libraries were prepared using the Illumina TruSeq DNA PCR-Free kit (Illumina, San Diego, CA) and sequenced as 150-bp paired-end reads on the NovaSeq 6000 platform. The mean genome coverage across all samples was 37.1× (minimum coverage being 30.1×). WGS and WES reads were processed separately and aligned to the human reference genome (GRCh38, GENCODE release 46)^22^ using DeepVariant^23^, applying the “WGS” and “WES” modes, respectively.

#### 4.2.2 Transcriptomics

Total RNA was isolated from whole blood obtained from 36 individuals, including 22 participants from the SPSS cohort and 14 from the EUCLIDS cohort. RNA-seq libraries were prepared using the Illumina TruSeq Stranded mRNA Library Prep Kit, following the manufacturer’s protocol. Sequencing was performed on an Illumina NovaSeq 6000 platform, generating 100 bp paired-end reads.

RNA-seq reads were aligned to the same human reference genome used for DNA analyses using STAR (version 2.7.7a)^24^. The alignment was performed in two-pass mode to enhance the detection of novel splice junctions.

Gene-level read counts were quantified using HTSeq (version 2.0.5)^25^, with the counting mode set to intersection-strict to include only reads fully contained within annotated gene features, and with count overlap enabled to maximize gene detection sensitivity.

#### 4.2.3 Proteomics

##### Mass Spectrometry Measurements

16 *µ*L of serum samples from the SPSS cohort or plasma samples from the EUCLIDS cohort were processed using the Mag-Net bead-based capturing protocol^26^. In short, the liquid biopsy was mixed 1:1 with binding buffer (100 mM Bis-Tris Propane, pH 6.3, 150 mM NaCl) containing protease inhibitor (Roche). All subsequent washing steps were performed on an Opentrons Flex robotics platform. MagReSyn^®^ strong anion exchange (SAX) beads (ReSyn Biosciences) were first equilibrated using equilibration/washing buffer (50 mM Bis Tris Propane, pH 6.3, 150 mM NaCl) and then combined in a 1 to 4 ratio with the liquid biopsy/binding buffer sample mix. The sample mix was incubated on the beads for 30 min at room temperature. The beads were then washed three times with equilibration/washing buffer. Proteins associated with the SAX beads were then solubilized and reduced in 50 mM Tris, pH 7.8 (after addition of TCEP) /2% SDS/10 mM Tris (2-carboxyethyl) phosphine (TCEP)/ 15 mM Chloroacetamide for 60 mins at 37 *^◦^*C in the dark. The samples were then processed further using aggregation capture by adding 100% acetonitrile to a final concentration greater that 70%. Samples were mixed and incubated for 30 min at room temperature. The beads were then washed three times in 95% acetonitrile and three times in 70% ethanol for 2 min each on the Opentrons Flex magnetic rack. Samples were then resuspended in 50 mM ammonium bicarbonate with containing 2 *µ*g Trypsin/LysC (Promega) per sample. After 16 hours of digestion at 37 *^◦^*C samples were quenched with formic acid to a final concentration of 0.2%. A quarter of the digested sample was then loaded onto C18 material containing Evotips together with iRTs (Biognosys) in a ratio of 1:40 v:v.

Peptides were separated by reversed-phase chromatography on an Evosep One liquid chromatography system (Evosep) using an Endurance column (EV1106 Endurance Column ReproSil-Pur C18, 1.9 *µ*m (Dr Maisch), 15 cm length x 150 *µ*m diameter) and a 30 samples per day method.

Peptides were injected into a Q Exactive HF-X high-resolution mass spectrometer (Thermo Fisher Scientific). Mass spectra were acquired using DIA-MS as described before^27–29^.

In short, one MS1 scan from 380 to 980 m/z was acquired at 60,000 resolution followed by 20 consecutive DIA segments (60 different segments in total) acquired at 15,000 resolution (AGC target 1 × 10e6 and auto for injection time) with fixed scan windows. Normalized collision energy was set to 28.

##### Data Analysis

DIA raw files were analyzed using Spectronaut v.19 in direct DIA mode against a UniProt database from January 2022. Carbamidomethylation was set as fixed modification, acetyl (protein N-term), deamidation (N) and oxidation (M) were set as variable modifications. Both peptide (MS1) and peptide fragment (MS2) data were used for peptide identification while quantification was based on MS1. FDR cut-off on the protein as well as on the peptide level was set to 0.01. Interference correction as well as a cross-run normalization was applied. Precursors were filtered based on their identification (Qvalue). For data analysis, proteomics datasets were filtered to exclude proteins with more than 50% missing values and to retain only proteins detected in both cohorts. Remaining missing values were imputed separately for each cohort using NAguideR^30^.

#### 4.2.4 Metabolomics

Metabolites were extracted from serum samples in the SPSS cohort and from plasma samples in the EUCLIDS cohort by methanol precipitation. Briefly, samples were thawed on ice for 30–60 min, and 20 *µ*L of sample was mixed with 180 *µ*L of 80% methanol (at room temperature). The mixture was vortexed for 15 s, incubated for 1 h at 4 *^◦^*C, and centrifuged at *>* 14,000 × *g* for 15 min (at room temperature).

Metabolite extracts were analyzed using flow-injection time-of-flight mass spectrometry (Agilent 6550 QTOF, negative ionization mode) as previously described^31^. This untargeted metabolomi approach scanned for ions in the 50–1,000 Da range, detecting more than 10,000 distinct mass-to-charge (*m/z*) features. High mass accuracy (∼1 mDa) and isotopic pattern analysis enabled the identification of several hundred metabolites across diverse chemical classes. Although structural isomers with identical elemental compositions could not be resolved due to the absence of chromatographic separation, elemental formulas and compound classes were determined with high confidence.

For each elemental formula, corresponding metabolites were annotated using the Human Metabolo Database (HMDB) 4.0^32^. Samples were analyzed sequentially (∼1,000/day) to minimize batch effects. Overall, the untargeted workflow quantified *2088* elemental formulas, corresponding to up to *1766* putative metabolites, including xenobiotics, drugs, and their metabolites.

### 4.3 Transcriptomic signature

To capture sepsis severity, we computed the quantitative sepsis response signature (SRSq) using SepstratifieR R package (https://github.com/jknightlab/SepstratifieR)^7^. As recommended for RNA-seq data, analyses were performed on log-transformed counts per million (log-CPM), with time to blood sampling (defined as the time interval in days between hospital admission and blood culture sampling) and batch effects regressed out using removeBatchEffect in limma^33^. We used the extended signature (19 genes) as all genes were detected and expressed above background (Supplementary Figure S1).

### 4.4 Multi-omics integration

#### 4.4.1 Multi-omics preprocessing

We integrated genomic, transcriptomic, proteomic and metabolomic layers into a shared latent space using the Multi-Omics Factor Analysis (MOFA) framework^10^. For this analysis, we applied layer-specific preprocessing and normalization separately within each cohort to harmonize feature distributions and enable comparability across datasets.

For the genomic layer, variants called by DeepVariant were normalized against the GRCh38 reference genome, followed by genotype-level filtering to remove low-depth calls (depth ≤ 20) and heterozygous variants with allelic imbalance (0.2 ≤ heterozygous allele balance ≤ 0.8). The resulting variant sets were harmonized and combined into a single cohort-level VCF, which was functionally annotated using Ensembl Variant Effect Predictor (VEP)^34^. Only variants predicted to have high or moderate functional impact were retained. For downstream integration, genotypes were encoded as binary features, with 1 indicating the presence of at least one alternative allele (heterozygous or homozygous alternative) and 0 indicating homozygosity for the reference allele, enabling modeling of the genomic layer in MOFA using a Bernoulli (binary) likelihood.

Because MOFA assumes Gaussian-distributed residuals for continuous data, appropriate normalization and variance stabilization were essential to satisfy model assumptions and enhance the interpretability of the inferred latent factors. RNA-seq counts were normalized using DESeq2^35^ with size-factor estimation, followed by variance-stabilizing transformation (VST). Transcriptomic features were subsequently adjusted for time to blood sampling. Proteomic and metabolomic data were similarly adjusted for time to blood sampling and transformed using log(*x* + 1) scaling to stabilize variance.

To explore potential regulatory variation beyond gene expression, we incorporated RNA splicing as an additional omics layer. Splicing quantification was performed using the FRASER2 pipeline^36^, which computes the ΔJaccard metric to quantify deviations in splice junction usage relative to a reference distribution. Only junctions overlapping annotated genes (excluding YRNAs) were retained to focus on biologically interpretable events. The resulting feature distributions were well suited for MOFA modeling and enabled the integration of splicing regulation alongside gene expression, protein abundance, and metabolite levels.

#### 4.4.2 Model optimization and feature selection

To identify shared molecular programs underlying pediatric sepsis across multiple omics layers, we applied the MOFA framework to the discovery SPSS cohort. MOFA is a well-established statistical framework for multi-omics data integration. Based on group factor analysis^37^, MOFA infers latent factors that explain sources of variation within and across data modalities. Model optimization focused on omics layers not constrained by detection limits (DNA, RNA, and RNA splicing) by selecting feature-set sizes that maximized the model’s total variance explained.

MOFA models were trained using the MOFA R package with 10 latent factors. We performed a grid search varying N from 0 to 2,500 (step 500), selecting the top-N most variable features within each data type. For each parameter combination, models were trained using three independent random seeds, and performance was evaluated using the mean total variance explained (*R*^2^) across runs, defined as the sum of per-view *R*^2^ across all omics layers. Default MOFA data, model, and training options were used. After selecting the final feature sets, we assessed latent factor stability by fitting 25 independent MOFA models initialized with different random seeds.

#### 4.4.3 Associations between latent factors and clinical or molecular variables

For each latent factor, associations with individual variables (demographic and clinical) were tested independently, with the statistical method chosen according to the variable type. Continuous variables were evaluated using Spearman’s rank correlation, whereas categorical variables were analyzed using non-parametric tests. Specifically, the Wilcoxon rank-sum test was applied for binary variables, and the Kruskal–Wallis test was used for multi-level categorical variables.

#### 4.4.4 Characterization of latent factors

In addition to assessing correlations with demographic, clinical, and molecular variables, we further characterized the MOFA-derived latent factors by examining their feature weights. Each feature weight represents the contribution of that feature to a given factor, with higher absolute values indicating stronger associations. The sign of the weight reflects the direction of the effect: positive weights correspond to features that show higher values in samples with positive factor scores, whereas negative weights correspond to features enriched in samples with negative factor scores.

To gain biological insight into the molecular processes captured by each factor, we performed gene set enrichment analysis (GSEA) on the transcriptomic and proteomic feature weights using the MOFA package built-in function. Enrichment analyses were conducted using the Reactome pathway database^38^. For each factor and molecular layer, we analyzed positively and negatively weighted features separately, allowing pathways associated with opposite directions of each latent factor to be evaluated independently.

In parallel, we estimated immune-cell composition from bulk RNA-seq data using CIBERSORT with the LM22 leukocyte signature matrix^39^. We correlated estimated immune-cell proportions with MOFA latent factor scores using Spearman’s rank correlation to assess whether latent molecular axes were associated with variation in immune-cell composition across sepsis samples.

### 4.5 Cross-cohort and cross-platform validation of the inflammatory axis

#### 4.5.1 Validation in an independent multi-omic dataset

To evaluate the generalizability of the multi-omics factors learned in the discovery cohort (SPSS), we performed external validation in the independent EUCLIDS cohort, comprising 22 pediatric patients with sepsis. The replication analysis aimed to determine whether the latent factors derived from the discovery cohort were conserved in the validation cohort. To ensure comparability, we restricted each omics layer to molecular features shared between cohorts (e.g., gene, protein, metabolite) and preprocessed the EUCLIDS data using the same pipeline and transformations applied to SPSS. Using these harmonized features, the EUCLIDS samples were projected onto the previously trained MOFA model to infer factor values for each individual, allowing direct comparison of latent representations between cohorts. Concretely, projection was performed by computing factor scores from the fixed, previously learned loadings via matrix inversion of the model weights. This approach estimates latent factor values for new individuals without retraining the model, thereby preserving the original latent structure.

#### 4.5.2 Validation in public transcriptomic datasets

To further evaluate whether the transcriptomic components of the discovered latent factors generalize to larger cohorts with controls available, we extended validation to publicly available whole-blood expression datasets.

Specifically, we validated the model using three pediatric whole-blood microarray expression profiles datasets from the Gene Expression Omnibus (GSE26440^40,41^ and GSE26378^41^, GSE13904^42^ All datasets consisted of whole-blood expression profiles collected within 24 hours of pediatric intensive care unit admission.

GSE26440 included 98 children with septic shock and 32 healthy controls, while GSE26378 comprised 82 children with septic shock and 21 controls. GSE13904 captured a larger clinical spectrum, with samples from 18 healthy controls, 22 children with systemic inflammatory response syndrome (SIRS), 32 with sepsis, and 67 with septic shock.

For each microarray dataset, probe-level expression data were preprocessed by applying background correction, normalization, and log transformation. Expression values from multiple probes mapping to the same gene were then averaged to obtain a single gene-level measurement. For each individual, SRSq was calculated from the processed output matrix using the 7-gene Davenport signature, as the complete gene set required for the extended signature was not consistently detected across samples. MOFA factor values were subsequently inferred for each sample by projecting centered and scaled expression data, limited to genes shared with the discovery cohort, onto the MOFA model using the inverse of the learned weight matrix. Factor scores were compared across clinical groups using Wilcoxon rank-sum tests for GSE26440 and GSE26378. For GSE13904, disease severity was analysed using cumulative link ordinal regression with clinical status encoded as an ordered outcome from healthy control to systemic inflammatory response syndrome, sepsis and septic shock, thereby testing whether factor scores tracked progression across the clinical spectrum.

## Acknowledgements

We thank the Health 2030 Genome Center for genomics and transcriptomics sequencing, and the PHRT Swiss Multi-Omics Center (SMOC) for proteomics and multi-omics data generation and preprocessing.

## Author contributions

**Study Design:** L.S, D.S.F, J.F

**Data management, participant selection:** R.M, A.A, V.Z, D.C, C.E.v.d.G.-d.J

**Data acquisition:** K.M, S.G, N.Z

**Data analysis:** M.A.O

**Drafting the manuscript:** M.A.O, A.S, B.R, J.F

**Critical review and revision of the manuscript:** All authors

## Data availability

The mass spectrometry raw files underlying the proteomic and metabolomic analyses will be deposited in a public repository such as MaSSIVE before publication, with accession numbers provided upon acceptance. Learned MOFA factor weights required to reproduce the main analyses, will be provided as supplementary tables. Public transcriptomic validation datasets used in this study are available from the Gene Expression Omnibus under accession numbers GSE26440, GSE26378 and GSE13904.

## Code availability

Custom scripts used for preprocessing, MOFA model training, factor projection, statistical analyses and figure generation will be made available in a public GitHub repository before publication, with a permanent archived version deposited in Zenodo.

