## Supplementary figures for "Multi-omics integration identifies a reproducible inflammatory host-response axis in pediatric sepsis"

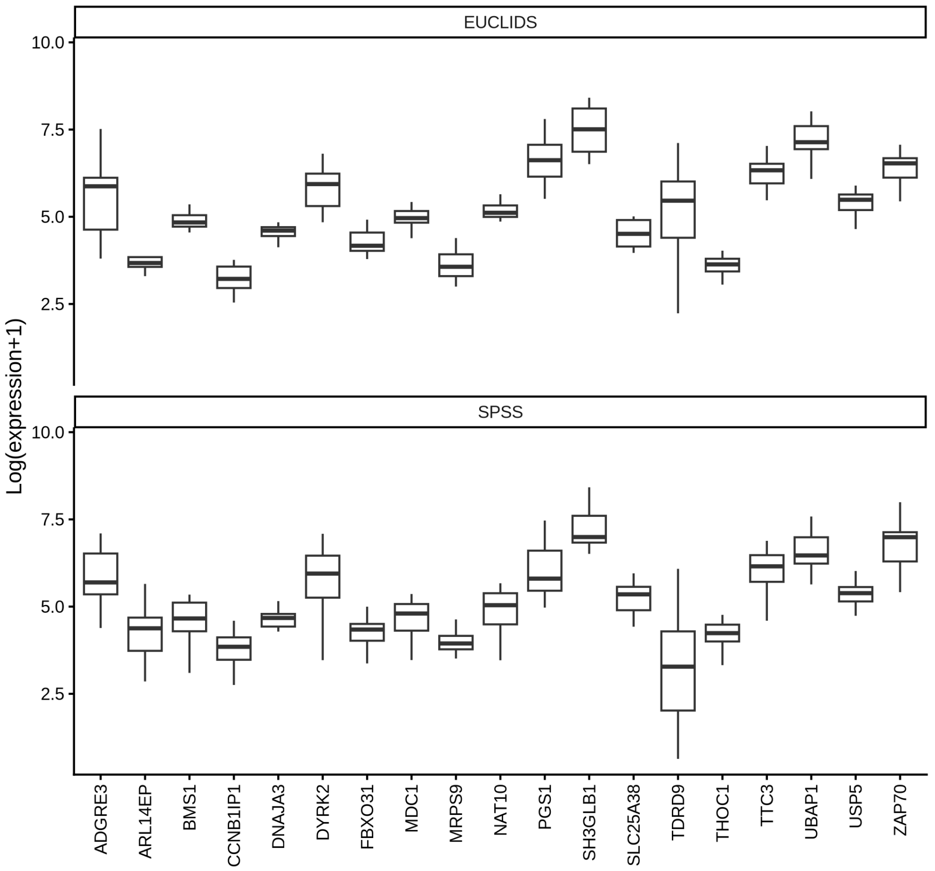


Figure S1: Distribution of log-transformed gene expression values for genes comprising the extended SRSq signature in two independent cohorts, EUCLIDS and SPSS. For each gene, the box indicates the interquartile range, the center line marks the median, and whiskers represent the spread of the data within each cohort.

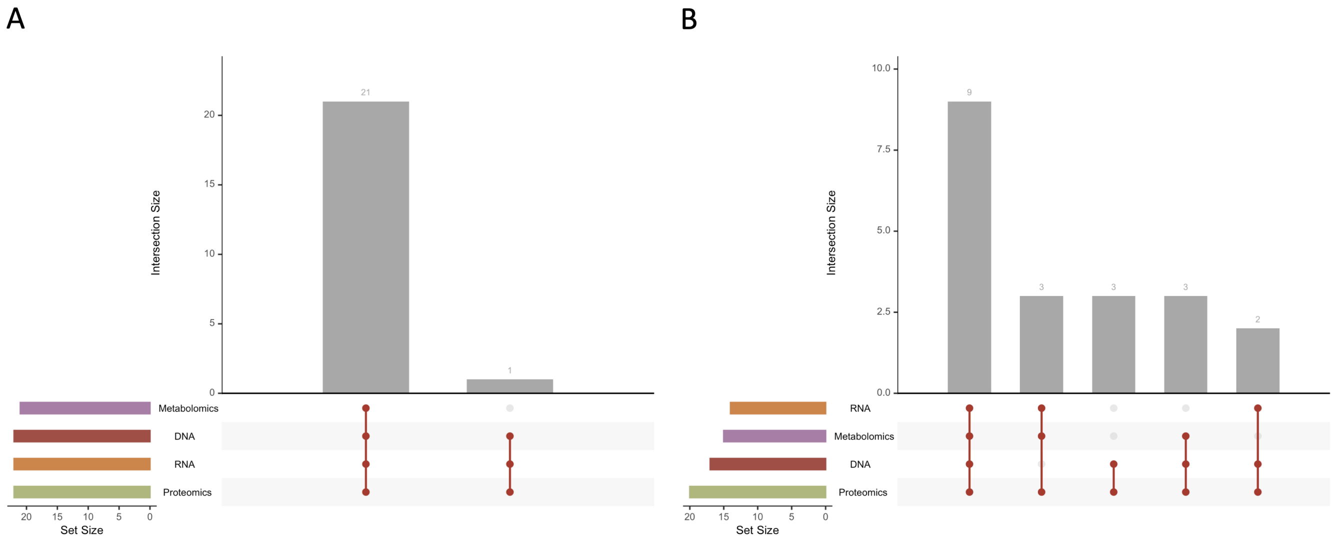


Figure S2: Data availability across omics layers in the SPSS and EUCLID cohorts. (A) UpSet plot depicting data availability for the SPSS cohort. (B) UpSet plot depicting data availability for the EUCLIDS cohort. Bars represent the number of participants with data available for each omic layer or combination of layers. Horizontal bars indicate the total number of participants with available data per omic type.

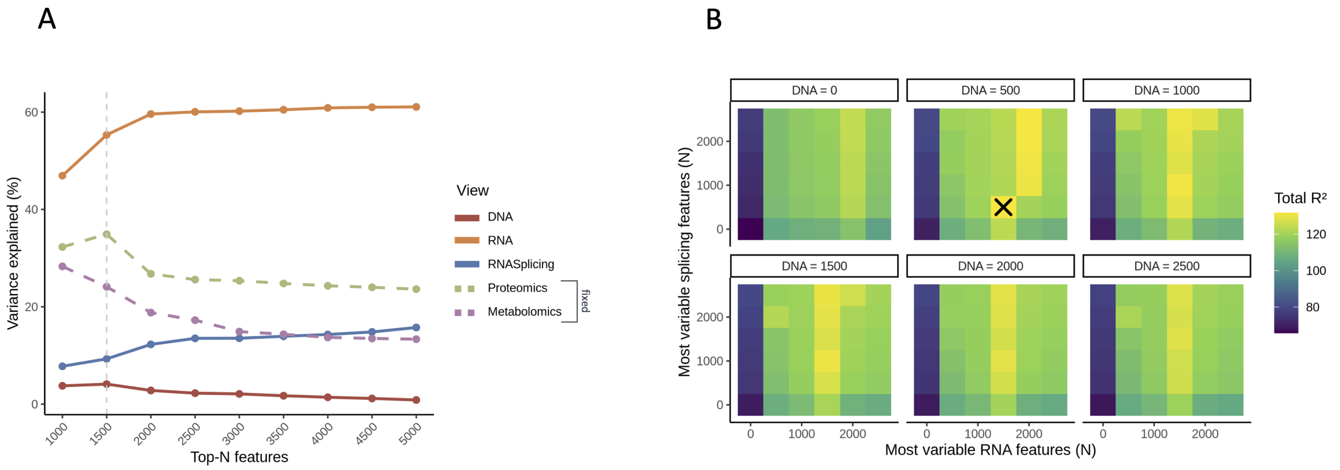


Figure S3: Optimization of feature selection in the MOFA model. (A) Fraction of variance explained by each omic layer as a function of the number of top-N most variable (DNA, RNA and RNA splicing) features included in the model. The dashed line indicates fixed number of features layers. (B) Grid search evaluating total variance explained ($R^2$) across combinations of top-N RNA and RNA splicing features for different DNA feature set sizes. The optimal configuration ($R^2$=131.71) (RNA = 1,500, RNA splicing = 500, DNA = 500) is indicated by an “X.”.

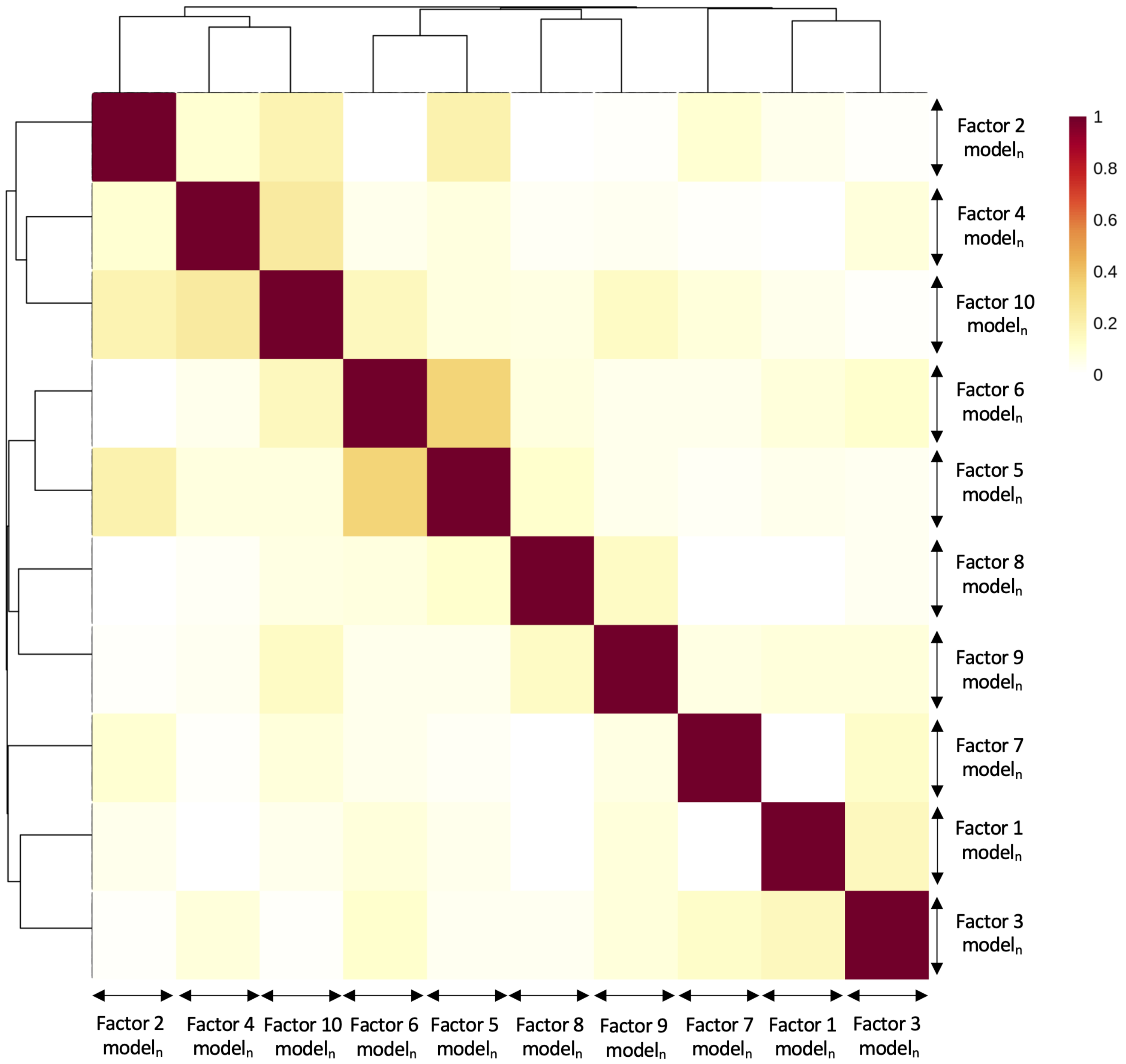


Figure S4: Reproducibility of MOFA latent dimensions across random initializations. Heatmap of pairwise correlations between latent dimensions inferred from 25 independent MOFA runs (different random seeds; 10 dimensions per run). Runs are indexed by n∈[0–25], indicating the different model initializations tested. The pronounced diagonal pattern and clustering of corresponding dimensions demonstrate stable recovery of the underlying latent structure across model initializations.

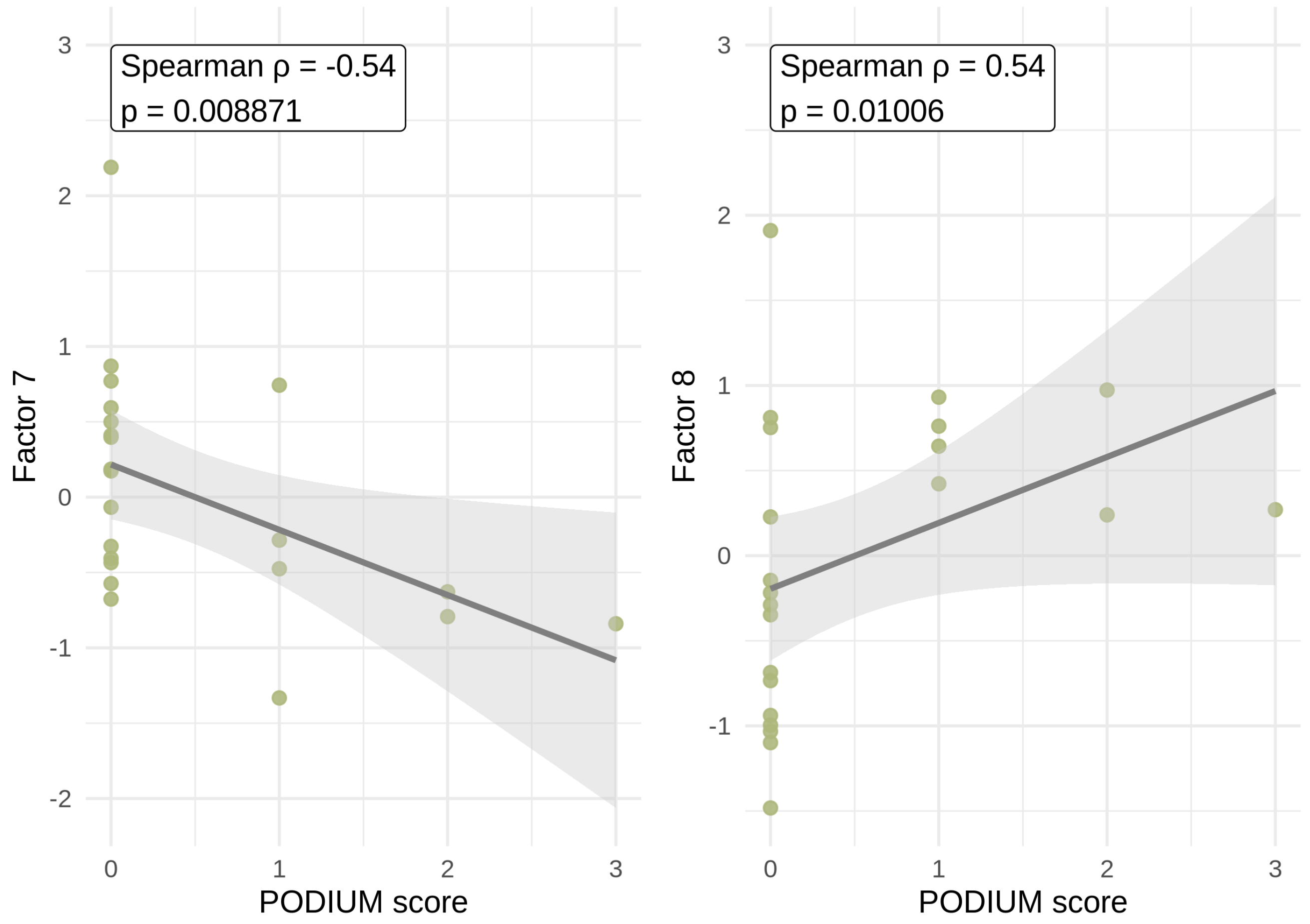


Figure S5: Association between the Pediatric Organ Dysfunction Information Update Mandate (PODIUM) score MOFA factor 7 (left) and factor 8 (right). Each point represents one individual. Lines indicate linear regression fits with shaded areas representing 95% confidence intervals.

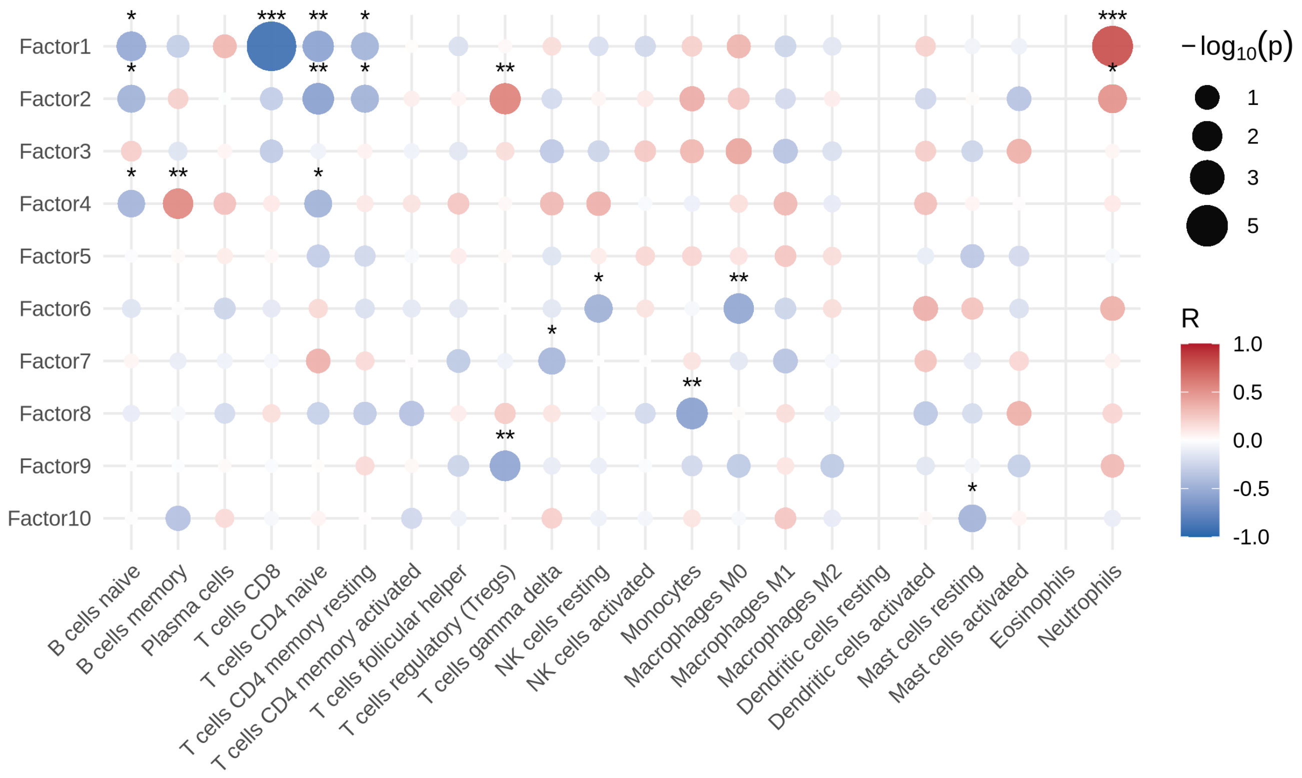


Figure S6: Associations between MOFA latent factors and inferred immune-cell composition in the SPSS discovery cohort. Bubble plot summarizing correlations between MOFA factor scores and relative immune-cell abundances estimated using CIBERSORT. Circle color indicates the direction and magnitude of the Spearman correlation coefficient (red: positive, blue: negative), and circle size reflects statistical significance ($-\log_{10}(P)$). Stars indicate significance levels ($*$, $**$, $***$ denote $P < 0.05$, 0.01, and 0.001, respectively).

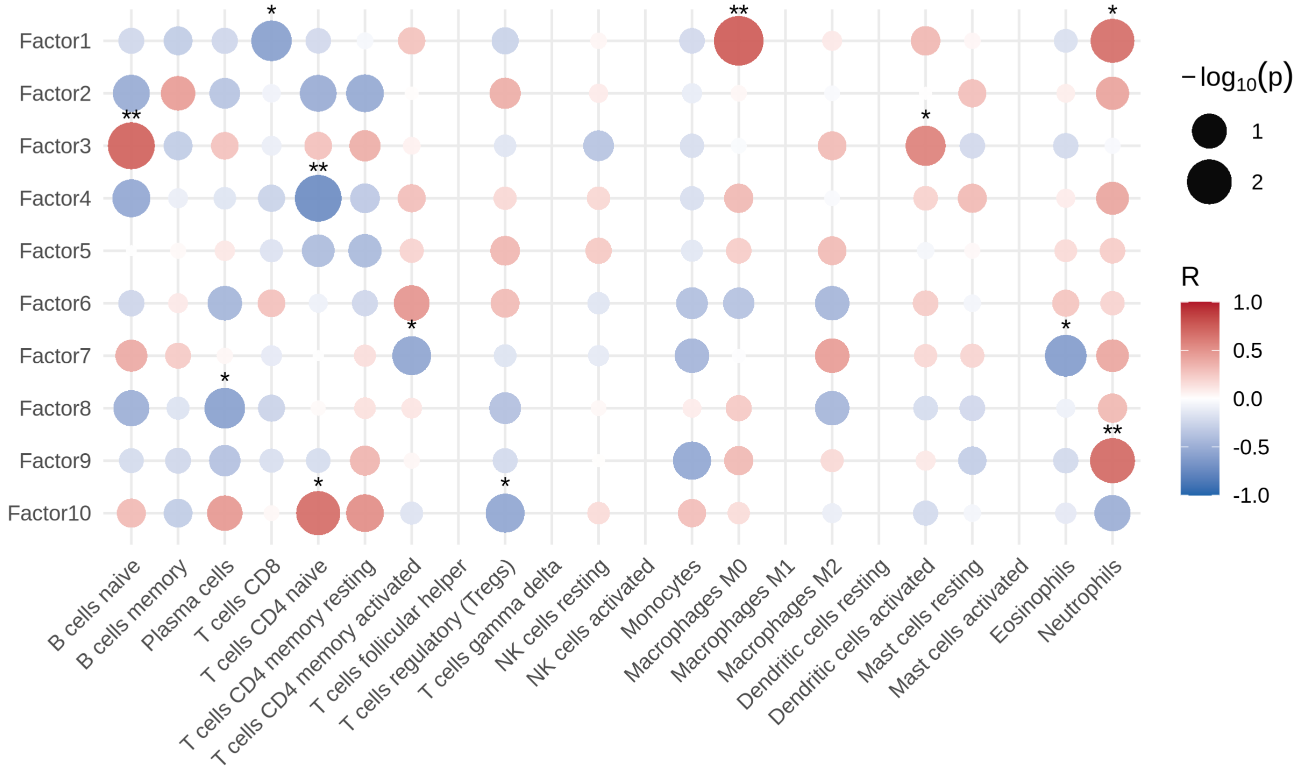


Figure S7: Associations between MOFA latent factors and inferred immune-cell composition in the EUCLIDS discovery cohort. Bubble plot summarizing correlations between MOFA factor scores and relative immune-cell abundances estimated using CIBERSORT. Circle color indicates the direction and magnitude of the Spearman correlation coefficient (red: positive, blue: negative), and circle size reflects statistical significance ($-\log_{10}(P)$). Stars indicate significance levels ($*$, $**$, $***$ denote $P < 0.05$, 0.01, and 0.001, respectively).

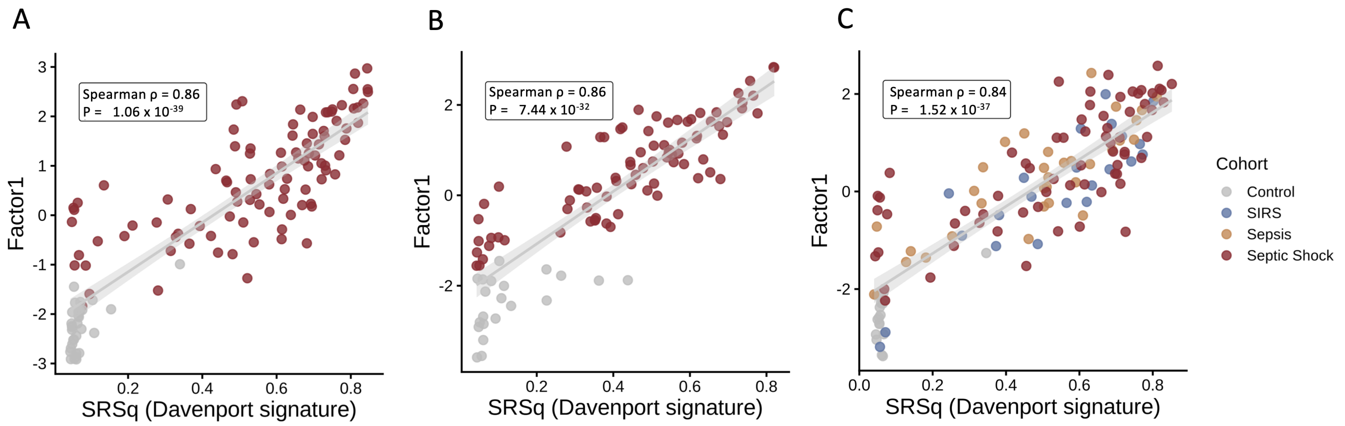


Figure S8: Correlation between Factor1 and the Davenport sepsis-response signature across cohorts. Scatter plots showing the association between Factor1 and the SRSq (Davenport signature) score in three independent transcriptomics datasets. Each point represents one sample. The solid line indicates the fitted linear regression, with shaded bands denoting the 95% confidence interval. Insets show the two-sided Spearman correlation coefficient ($\rho$) and corresponding $P$ value for each panel.

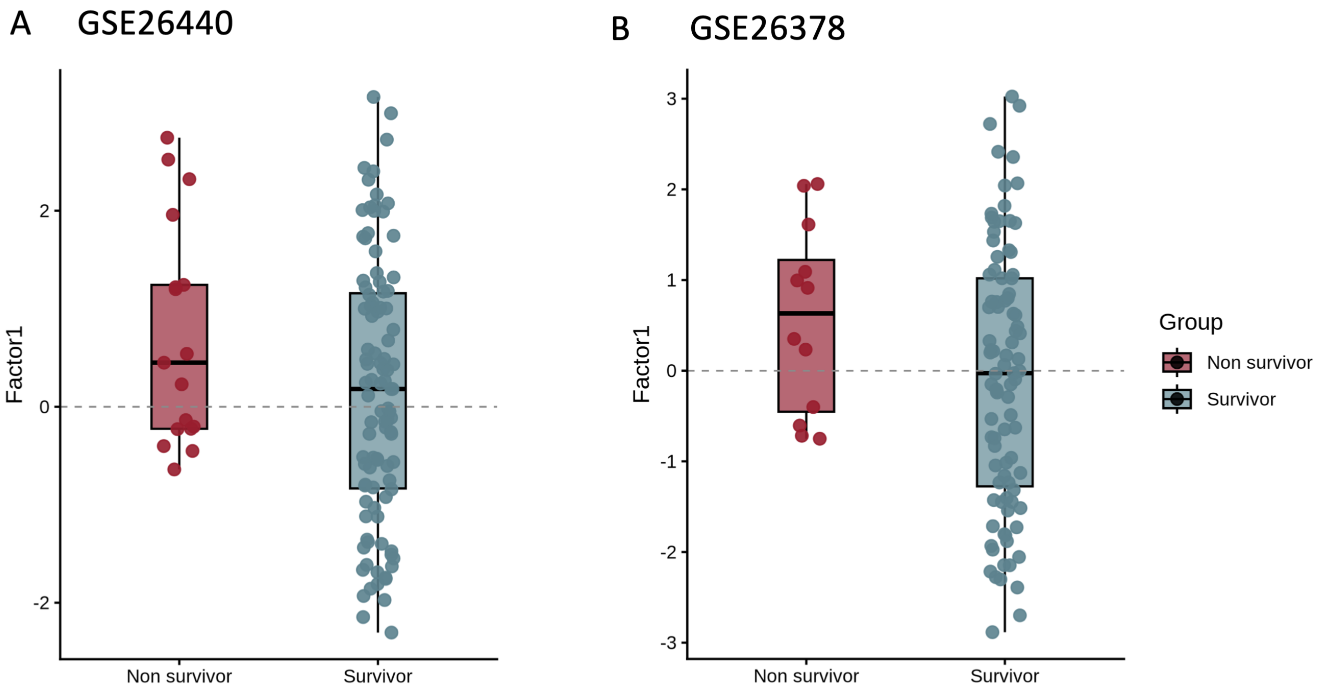


Figure S9: Distribution of Factor 1 scores in two independent GEO pediatric sepsis cohorts (GSE26440 and GSE26378) comparing non-survivors (red) and survivors (blue).

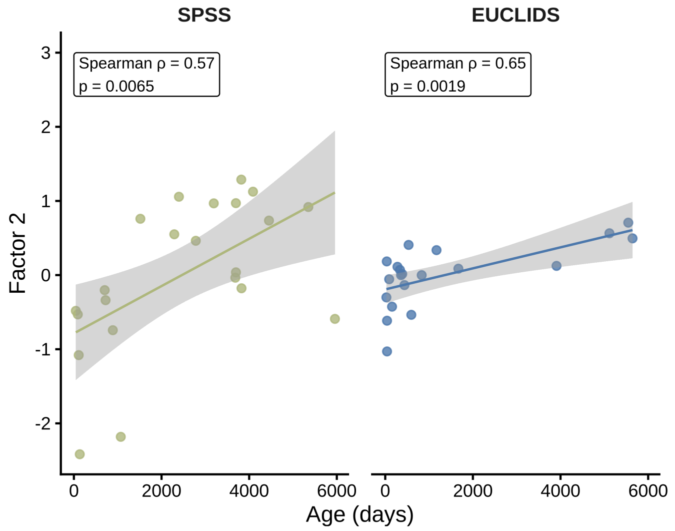


Figure S10: Association between MOFA factor 2 and age at sampling across the SPSS (left) and EUCLIDS (right) cohorts. Each point represents one individual. Lines indicate linear regression fits with shaded areas representing 95% confidence intervals.

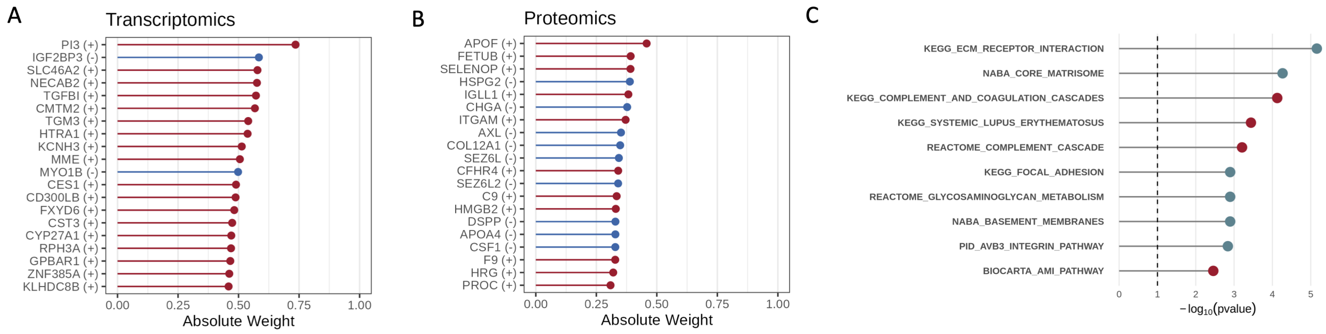


Figure S11: Molecular features contributing to MOFA factor 2 and pathway enrichment of proteomic loadings. (A)Top transcriptomic features contributing to factor 2, ranked by absolute loading weight. (B) Top proteomic features contributing to factor 2, ranked by absolute loading weight. Several high-weight proteins are involved in extracellular matrix organisation, complement regulation and coagulation pathways. (C) Gene set enrichment analysis (GSEA) of proteomic loadings using MSigDB C2 canonical pathways. Points represent pathway enrichment significance expressed as $\log_{10}$(p value). Colors indicate direction of association with factor 2 (red, positively associated/upregulated; blue, negatively associated/downregulated).
